# A Machine Learning Model of Microscopic Agglutination Test for Diagnosis of Leptospirosis

**DOI:** 10.1101/2020.12.08.410712

**Authors:** Yuji Oyamada, Ryo Ozuru, Toshiyuki Masuzawa, Satoshi Miyahara, Yasuhiko Nikaido, Fumiko Obata, Mitsumasa Saito, Sharon Yvette Angelina M. Villanueva, Jun Fujii

## Abstract

Leptospirosis is a zoonosis caused by the pathogenic bacterium *Leptospira*. The Microscopic Agglutination Test (MAT) is widely used as the gold standard for diagnosis of leptospirosis. In this method, diluted patient serum is mixed with serotype-determined Leptospiras, and the presence or absence of aggregation is determined under a dark-field microscope to calculate the antibody titer. Problems of the current MAT method are 1) a requirement of examining many specimens per sample, and 2) a need of distinguishing contaminants from true aggregates to accurately identify positivity. Therefore, increasing efficiency and accuracy are the key to refine MAT. It is possible to achieve efficiency and standardize accuracy at the same time by automating the decision making process. In this study, we built an automatic identification algorithm of MAT using a machine learning method to determine aggregation within microscopic images. The machine learned the features from 316 positive and 230 negative MAT images created with sera of Leptospira- infected (positive) and non-infected (negative) hamsters, respectively. In addition to the acquired original images, wavelet-transformed images were also considered as features. We utilized a support vector machine (SVM) as a proposed decision method. We validated the trained SVMs with 210 positive and 154 negative images. When the features were obtained from original or wavelet-transformed images, all negative images were misjudged as positive, and the classification performance was very low with sensitivity of 1 and specificity of 0. In contrast, when the histograms of wavelet coefficients were used as features, the performance was greatly improved with sensitivity of 0.99 and specificity of 0.99. We confirmed that the current algorithm judges the positive or negative of agglutinations in MAT images and gives the further possibility of automatizing MAT procedure.

## Introduction

Leptospirosis, an infectious disease caused by the pathogenic species of *Leptospira*, is one of the most widespread zoonoses in the world. The World Health Organization (WHO) estimates one million leptospirosis cases and 58,900 deaths worldwide each year, of which more than 70% occurring in the tropical regions of the world ^1^. Nonspecific and diverse clinical manifestations make clinical diagnosis difficult, and it is easily misdiagnosed with many other diseases in the tropics, such as dengue fever, malaria, and scrub typhus ^2^.

Microscopic agglutination test (MAT) is considered as the standard test for serological diagnosis of leptospirosis ^3^. Leptospires have over 250 serovars ^4^ and MAT is usually used to diagnose patients based on the *Leptospira* serotypes that infect humans or animals. The principle of MAT is simple, it consists of mixing the serially diluted test serum with a culture of leptospires and then evaluating the degree of agglutination due to immunoreaction using a dark-field microscope ^2^. The highest serum dilution agglutinating 50% or more of the leptospires is considered to be the antibody titer. However, the procedures involved in MAT, especially judging the results (i.e., whether positive or negative) requires highly trained personnel that making it difficult to adopt as a general test ^5^. Furthermore, the liquid handling such as transferring all the samples from each well of multi plates onto slide grasses is complicated and takes time. Although the International Leptospirosis Society has been implementing the International Proficiency Testing Scheme for MAT for several years now ^6^, worldwide standardization of MAT is yet to be achieved. This is partially because not only devices used for MAT such as dark-field microscopes, objective lenses, illuminations and cameras, but the testing condition (the dilution range of serum, incubation time and magnification of an objective lens used) of MAT is diverse at the various laboratories.

This kind of diagnosis is solved as binary classification whether each data is categorized either as positive or negative. Binary classification consists of training and test steps. Figure 1 shows an example of 2-dimensional non-linearly separable and inseparable data. Figure 1A shows a good data where positive and negative data are separably distributed. The training step makes several potential boundaries (curves in Fig. 1B) and selects the boundary that separates the training data with the highest classification score (the green curve in Fig. 1B). As shown in the figure, the selected boundary perfectly separates the negative and positive data. The test step evaluates the selected boundary with test data (Fig. 1C). As shown in the figure, the selected boundary both negative and positive data are perfectly separated, which means the trained classifier is good. The success of binary classification heavily depends on the distribution of data. Figure 1D-1F shows how badly distributed data worsen the performance of binary classification. Contrast to the good case, Fig. 1D shows a bad data where positive and negative data are inseparably distributed. None of the potential boundaries cannot perfectly separate even the training data (Fig. 1E and 1F).

**Fig 1.**
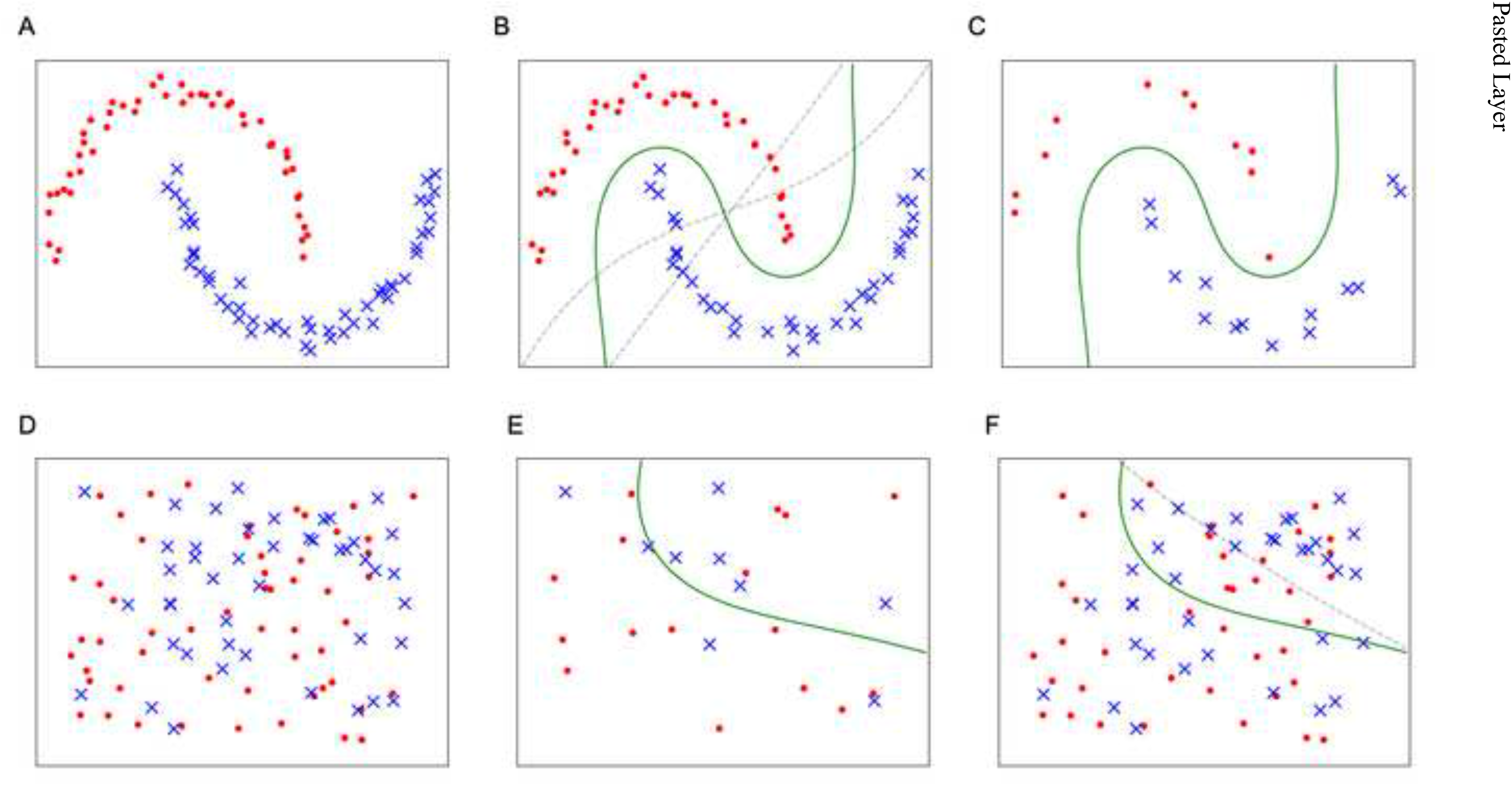
An example of 2-dimensional non-linearly separable and non-separable data for binary classification. Red circle and blue cross symbols represent negative and positive data. Curves represent boundaries obtained from training data and green one represents the best boundary. (A) A good training data. (B) Potential boundaries obtained from (A). (C) The best boundary obtained from (A). (D) A bad training data. (E) Potential boundaries obtained from (B). (F) The best boundary obtained from (B).

For binary classification on images, we use image features instead of raw images because raw images are too complex and redundant, which is potentially badly distributed data. An image feature, extracted from an image, encodes interesting information of the image into a sort of numerical values such as vector and matrix ^7^. Image features appropriate for binary classification are sensitive to class differences but not to other factors, which means each negative and positive data are distributed separately. Therefore, it takes an important role to use or design appropriate image features for the target problem.

In this paper, we developed a model to verify the possibility of applying machine learning technique to MAT. This is the first report on machine learning techniques for MAT diagnosis that uses a binary classification where each MAT image is taken either as agglutination positive or negative. This study aims to: (1) establish a machine learning based method for reading MAT results to aid in serodiagnosis; (2) design a MAT image feature that is appropriate for binary classification based on general machine learning and image processing techniques; and (3) conduct a evaluation test with MAT images obtained from animal experiments. This study will be the first step of our ultimate goal that is to fully automate the MAT procedure.

## Materials and Methods

This section describes the proposed method that uses machine learning techniques for MAT image classification that takes each MAT image either as agglutination positive or negative. Our idea is to follow a typical machine learning based image classification pipeline so that we can efficiently and effectively classify MAT images. The key contribution of this paper is to design an appropriate MAT image feature making binary classification feasible. The design of the image feature is based on how expert examiners evaluate MAT images. They consider how agglutination occupies a MAT image and therefore the proposed method extracts occupancy like information from a MAT image. The remainder of this section first explains how datasets were collected for this research and then the major steps, pre- processing, feature extraction, and binary classification issues were described.

### Animal Ethics Statement

Animal experiments were reviewed and approved by the Ethics Committee on Animal Experiments at the University of Occupational and Environmental Health, Japan (Permit Number: AE15-019). The experiments were carried out under the conditions indicated in the Regulations for Animal Experiments of the said University and Law 105 and Notification 6 of the Government of Japan.

### Preparation of image datasets

#### Animal infection experiment and acquisition of MAT images

In this study, a total of three golden syrian hamsters, male, 3-week-old were used (purchased from Japan SLC, Inc., Shizuoka, Japan). Serum from a hamster subcutaneously infected with 10^4^ Leptospira interrogans serovar Manilae at day 7 post infection was used as a MAT positive sample, and sera from other two hamsters at day 0 post infection was used as a MAT negative. MAT was performed according to the standard manual (Terpstra et al., 2003). In brief, each serum was primarily diluted 50-fold and then serially diluted (2-fold) until 25,600-fold dilution. Leptospires (1 × 10^8^) was added to each diluted serum and incubated at 30°C for 2-4 hours. Samples from each well were transferred to slide glasses and covered with coverslips. Each sample was observed under a dark-field microscope (OPTIPHOT, Nikon). Ten images from 20× or 40× objective lens fields per slide were obtained with a CCD camera (DP21, Olympus).

#### Pre-processing

In this step, training and test datasets were prepared from the original image datasets as shown in Fig. 2. As it was mentioned earlier, the proposed method utilizes the occupancy of agglutination area in a MAT image as an image feature. This assumes that all the images are taken under the same condition such as lighting condition and meter-resolution scale.

**Fig 2.**
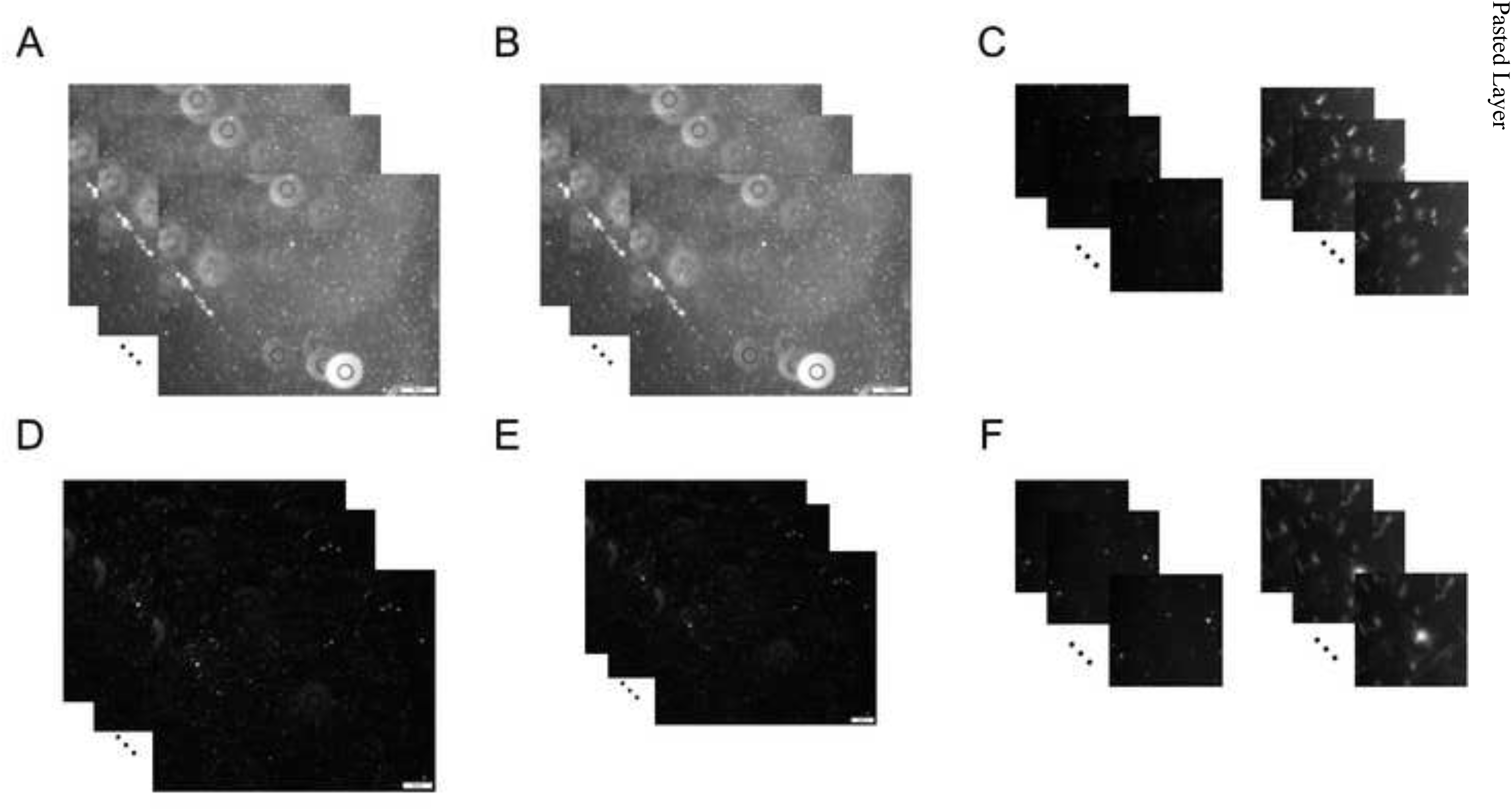
The data follow of the Pre-Processing. (A) Raw MAT images of Negative data. (B) Scale normalized images of Negative data. (C) Extracted patches of Negative data. (D) Raw MAT images of Positive data. (E) Scale normalized images of Positive data. (F) Extracted patches of Positive data.

Furthermore, images are supposed to contain enough resolution to measure the occupancy of areas with agglutination but not too large in order not to overwhelm the computer. To limit sizes of images suitable to analyze, all images were resized into smaller patches prior to the feature extraction.

#### Scale normalization

The scale normalization is done in order to make all the images in the dataset have the same micrometer-resolution scale. The original dataset is made of a series of negative images captured under 20X objective lens (50 micrometer scale) and of positive images with 40X objective lens (20 micrometer scale) (Fig. 3A and 3B). They contain the same resolution; however, the pixels applied to 50 micrometer in the negative images are 290, whereas to 20 micrometer in the positive images are 232. Based on the micrometer-resolution scale indicator shown in each image, this preprocessing resizes them to have the same micrometer-pixel scale (Fig. 3C and 3D).

**Fig 3.**
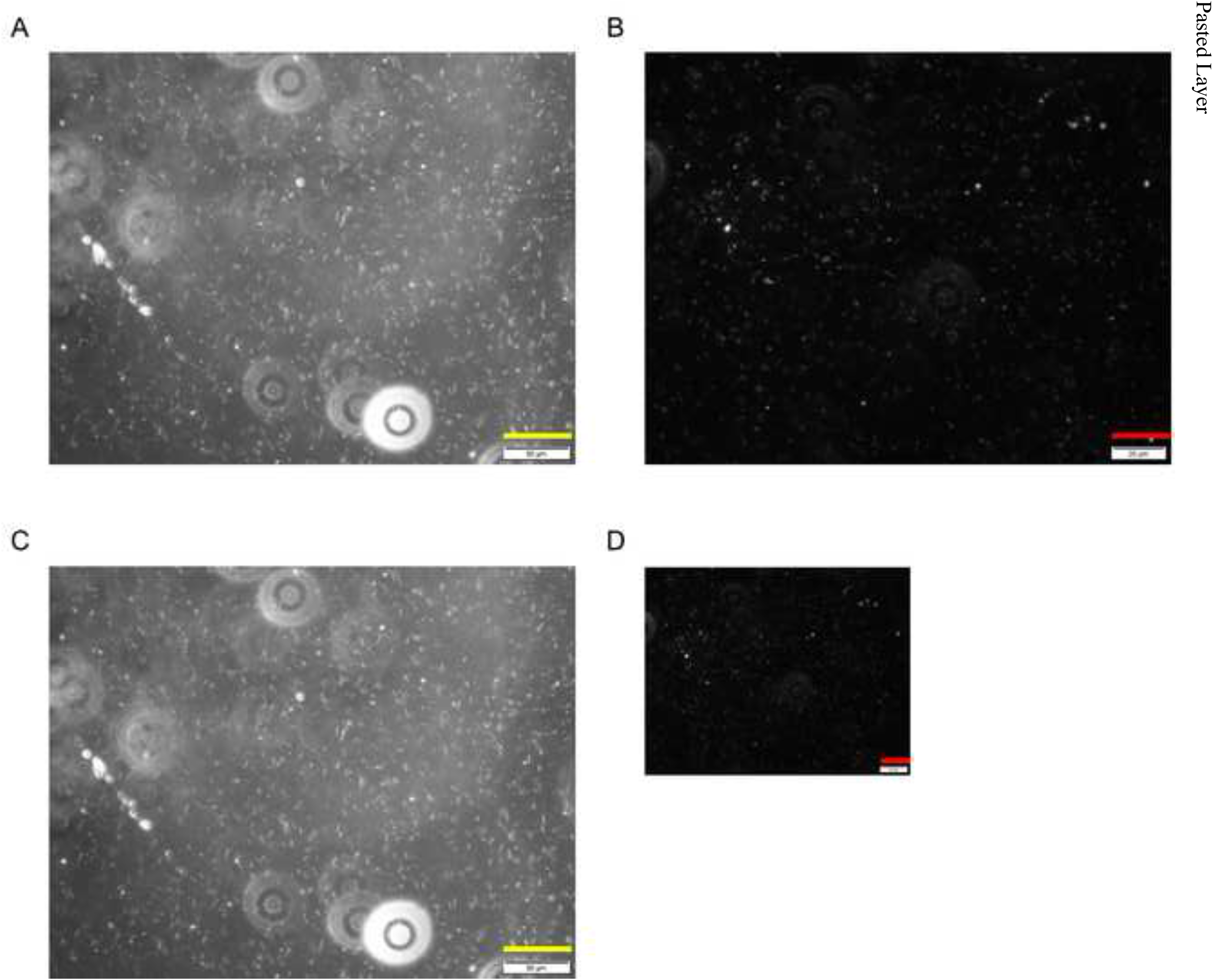
An example of the scale normalization. Yellow and red lines represent 50 and 20 micrometers respectively. (A) A raw MAT image of Negative data. (B) The scale normalized (A). (C)A raw MAT image of Positive data. (D) The scale normalized (C).

### Image patching

In the next step, each image was cut into smaller patches. Each patch must be large enough to count the amount of agglutination and also must avoid too large resolution to fulfill the minimum requirement of computers. In the succeeding experiments, different patch sizes were compared and the effect of the patch size difference was validated.

### Patch-level standardization

The last step of the pre-processing was to apply data standardization, a frequently used data normalization method, on each patch. This process was applied to satisfy an assumption by the later classification step that all feature vectors are centered around zero and have similar variance. For instance, if the data acquisition process affects the brightness of observed pictures, then the standardization is applied to cancel those unwilling effects. Suppose we have a feature vector *x* ∈ *R*^*d*^ representing an image patch. Data standardization normalizes the feature vector such that the normalized vector has zero mean and unit variance as 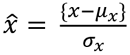, where *μ*_*x*_ and *σ*_*x*_ denote the mean and standard deviation of a set of *x*.

### Image features

Here, the design of the image feature that takes the key role of image classification was described (Fig. 4). There was still difficulty in the pre-processed MAT image patches such as images that might contain non-agglutination objects such as dust that has strong reflection. Consequently, non-agglutination objects from the pre-processed MAT image patches are excluded and the occupancy of the agglutination area is measured. Specifically, the designed image feature is the histogram of multi-level wavelet coefficients of a MAT image patch. The multi-level wavelet transform (Mallat, 1989) was used to extract objects on MAT image patches that have similar size to agglutination. By using the histogram of its coefficients, the behavior of the coefficients can be efficiently represented. In the remainder of this part, raw images (Image), multi-level wavelet coefficients (Wavelet), and the histogram of multi-level wavelet coefficients (*HoW*) will be explained step by step.

**Fig 4.**
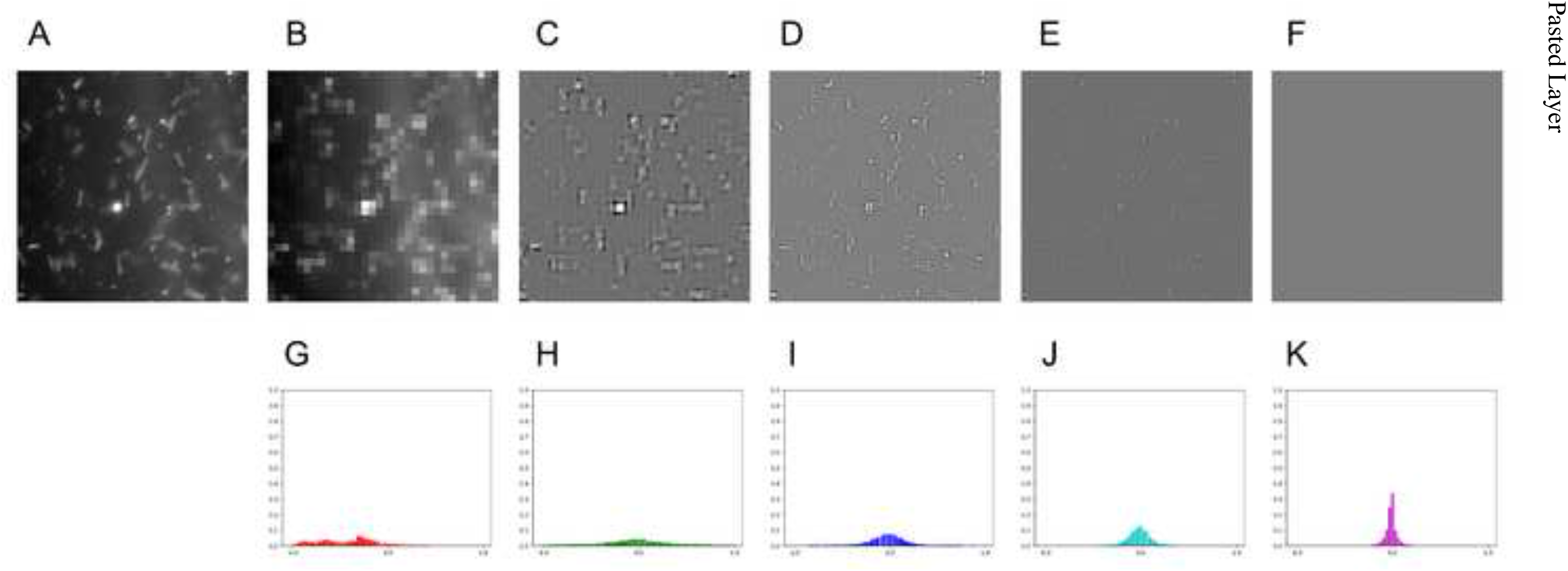
An image and its image features. (A) An image. (B)-(F) The 0-th to 4-th level wavelet coefficients of (A) (*Wavelet-0, …, Wavelet-4*). (G)-(K) The histogram of the 0-th to 4-th level wavelet coefficients of (A) (*HoW- 0, …, HoW-4*).

#### Raw images (*Image*)

One of the simplest image features is raw image itself.

A gray-scale image is represented as a 2D matrix *I* ∈ *R*^*X* × *Y*^, where *X* and *Y* denote the image width and height. A pixel intensity at pixel(*x*, *y*) is represented as *I*_*x*,*y*_ and the value is usually expressed by an eight-bit integer, 0, 1, 2, …, 255. When raw images are used as an image feature, comparison between two images was done by comparing their corresponding pixel intensities. For instance, mean square error between two images *I* and *I*^′^ is defined as

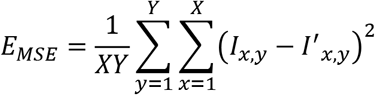

Raw images are easy to use and require less computer power. However, the pixel-wise comparison is inappropriate to consider the occupancy of the agglutination areas.

### Multi-level wavelet coefficients (*Wavelet*)

To extract agglutination of a certain size, multi-level wavelet transformation that is usually used to decompose images into base signals in different scales was utilized.

The idea of multi-level wavelet transformation (Mallat, 1989), which decomposes an image to a combination of a base signal with different directions and resolutions, is similar to the Fourier transformation. Applying multi-level wavelet transformation to a MAT image patch, any objects on the patch are separately extracted and stored into coefficients at different scales as larger coefficients. The lowest scale coefficients *W*_0_ ∈ *R*^*X*_*l*_×*Y*_*l*_^ is the mean image of the patch while the other coefficients {*W*_*l*_ ∈ *R*^*X*_*l*_×*Y*_*l*_^ | *l* = 1, . . . , *L*} contain large values where image objects of specific size exist at the location, where *X*_*l*_ and *Y*_*l*_ denote the width and height of l-th level as

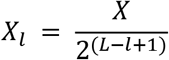

and

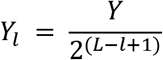

Namely, lower-level coefficients have the information of smaller image objects and larger level coefficients have one of larger image objects. In other words, lower-level coefficients correspond to coarser image components such as rough shape objects and larger level coefficients to finer components.

Figure 4A-4F show an image and its multi-level wavelet coefficients with Haar wavelet as base signal and 4 levels. As shown in the figure, image objects of different sizes are extracted in different levels, coarser features are in lower level and finer in higher. In these figures, zero coefficients are depicted as gray pixels and strong features are depicted as white and black pixels. Trained MAT examiners confirmed that *Leptospira-*like objects showed brighter pixels in levels 2 and 3 but not in the other levels.

Multi-level wavelet transformation to MAT image patches was applied and decided their coefficients of a level might be a candidate of the image feature. Benefit of this feature is that we can exclude non-agglutination objects based on their size. The drawback is the same as the *Image* feature that *Wavelet* feature also applies pixel-wise comparison.

#### Histogram of Multi-level wavelet coefficients (*HoW*)

To represent the occupancy of the agglutination areas in a MAT image patch, a normalized histogram of multi-level wavelet coefficients per level with a sum of 1 was constructed. The shape of such histograms represents the occupancy of the agglutination areas.

Constructing the histogram of wavelet coefficients is equivalent to count agglutination-like objects and therefore it is appropriate to measure the occupancy of the agglutination areas. Multi-level wavelet transformation was first applied and the histogram of the coefficients per level was constructed. Next, the histogram with a sum of 1 was normalized. By normalizing the histogram, histograms of different image resolutions are comparable. In total, we have a set of *L* + 1 histograms as *H*_*l*_ ∈ *R*^*B*^ | *l* = 0, . . . , *L*} where *H*_*l*_ denotes the normalized histogram of *l*-th level wavelet coefficients and *B* denotes the number of histogram bins. Benefits of this feature are its robustness against the existence of non-agglutination objects.

Figure 4G-4K show *HoW*s at different levels of an image. As shown in the figure, histograms at different levels have different sharpness. Since each level of wavelet coefficients has zero at most of the pixels, their histogram has their peak at the center. When the patch contains image objects of a specific size, their corresponding histograms have gentle peaks.

### Image feature classification

Here, we explain the details of binary classification. The proposed method utilizes the Support Vector Machine (SVM) (Boser et al. 1992), which is one of the supervised learning models for classification problems and has been well-used in practical situations because of its generalization performance against unknown data. Moreover, grid search hyper- parameter optimization and K-fold cross validation were combined in order to obtain better training effects.

#### SVMs

The proposed method utilizes Support Vector Machines (SVMs)^8^, which is one of well-used supervised learning models for solving classification problems.

Suppose we have a set of *N*training data *D* = {(*x*_*i*_, *y*_*i*_) | *x*_*i*_ ∈ *R*^*M*^, *y*_*i*_ ∈ {−1, 1}| *i* = 1, …, *N*}, where *x*_*i*_ denotes feature vector and *y*_*i*_ denotes its label. The label *y*_*i*_ = −1 indicates*i*-th feature is negative data and *y*_*i*_ = 1 indicates it is positive data. In our case, *x*_*i*_ is a MAT image feature, and negative and positive data means non-agglutinated and agglutinated data, respectively. Given the set of training data *D*, an SVM finds a boundary that maximizes its margin, which is the largest distance to the nearest data of both classes.

Using kernel trick, SVMs accomplish even non-linear classification. Using a kernel function, An SVM alters original data *x*_*i*_ to higher dimensional space, and linearly separates the calculated data in the projected domain. The kernel function decides the complexity of boundary shape.

Typical kernel functions for non-linear classifications are polynomial functions

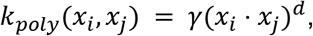

where *γ* denotes the scale factor and *d* the dimensionality of the polynomial and radial basis functions

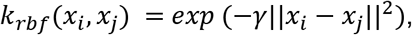

where *γ* denotes the non-negative scale factor.

#### Grid Search Hyper-parameter tuning

The proposed method utilizes grid search for tuning hyperparameters of SVMs.

The hyperparameters of SVMs are parameters of kernel function, e.g., *γ* of polynomial and radial basis functions, and general SVM parameter *C*. Grid search is one of hyperparameter tuning methods that exhaustively considers all potential combinations of hyperparameters.

#### K-fold cross validation

The proposed method applies K-fold cross validation to maximize training effects.

One of key concerns about supervised learning models is its generalization performance against unseen data. SVM training is similar to how we tune all parameters for the given training data but not for unseen data. So, there is no guarantee that the trained SVM performs well against unseen data. Situations that a trained supervised model performs well to training data but not to unseen data is called overfitting.

K-fold cross validation is a well-used practical solution to avoid such overfitting. Figure 5 shows the concept of K-fold cross validation. All data is split into either training or test datasets. The training dataset is further divided into *K* group, each of which is called a fold. For each fold, an SVM is trained with the remaining *K* − 1 folds and is validated with the fold. The validation errors of all folds are averaged to measure generalization performance of the model while cancelling overfitting for specific data. After the training, the trained SVM is further validated with the test dataset.

**Fig 5.**
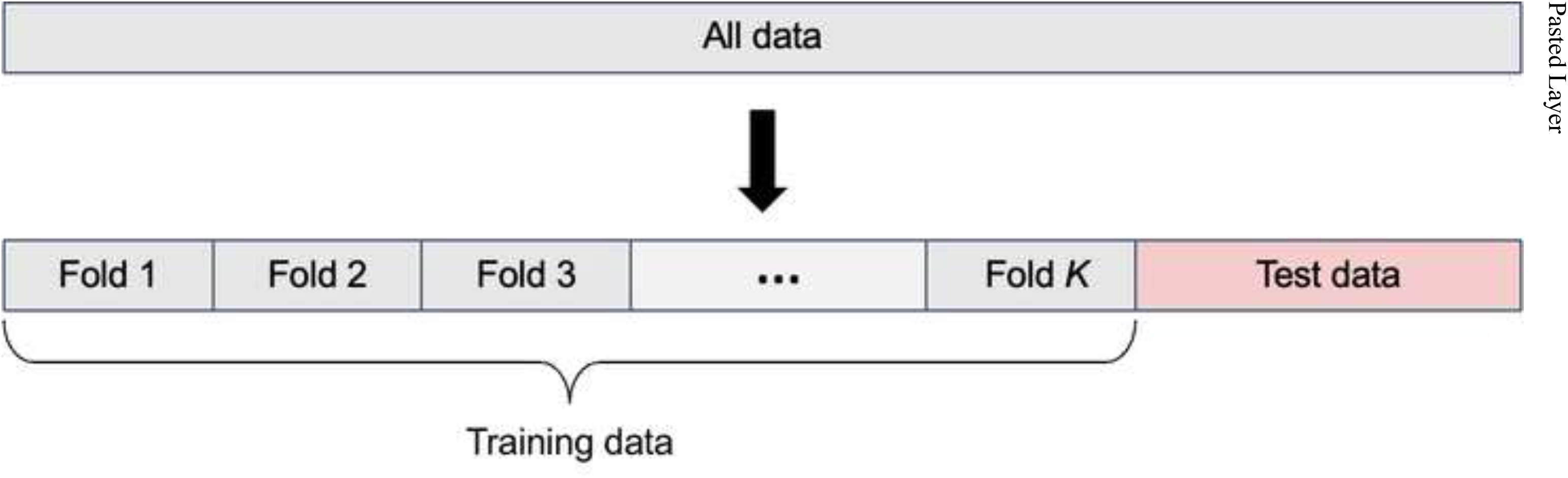
The concept of K-fold cross validation.

## Results

We conducted three evaluations to validate the potential of machine learning and image processing techniques on MAT: (1) the elapsed time of the feature extraction process, (2) qualitative evaluation of the MAT image feature classification, and (3) quantitative evaluation of the MAT image feature classification.

### Experimental conditions

Here, we describe the experimental conditions. We conducted all the experiments on a computer with an 8-core central processing unit (CPU) (Intel(R) Xeon(R) CPU E5-2620 v4) and 128 GB random access memory (RAM). All the data is stored on a hard disk drive (HDD). We implemented the proposed method using Python with standard packages such as numpy and scikit-learn and PyWavelet for Multi-level Wavelet Transformation ^9^.

Table 1 and 2 show the data properties. We have 295 images in total, which consist of 32 negative and 263 positive images. All images are in a resolution of 2448 x 1920. The micrometer-pixel ratio is 5.8 and 11.6 in negative and positive images, respectively. We resized all positive data so that the micrometer-pixel ratio is consistent across the classes.

**Table 1.** The specification of raw MAT images.

| | #Data | Resolution [pixels] | Scale [$\mu\text{m}/\text{pixel}$] |
| --- | --- | --- | --- |
| Negative | 32 | 2448x1920 | 5.8 |
| Positive | 263 | 2448x1920 | 11.6 |

**Table 2.**
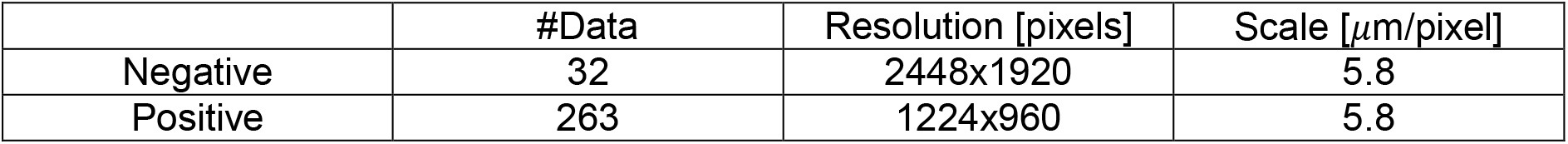
The specification of scale normalized MAT images.

| | #Data | Resolution [pixels] | Scale [$\mu\text{m}/\text{pixel}$] |
| --- | --- | --- | --- |
| Negative | 32 | 2448x1920 | 5.8 |
| Positive | 263 | 1224x960 | 5.8 |

In these experiments, we test two patch sizes 256 x 256 and 512 x 512. The number of patches for each patch size is shown in Table 3.

**Table 3.** The specification of scale normalized MAT images.

|  | 256x256 | 512x512 |
| --- | --- | --- |
| Negative | 1680 | 360 |
| Positive | 2688 | 448 |
| Total | 4368 | 808 |

For image features, *L*, the level of multi-level wavelet transformation, is set to 4, whereas *B*, the number of histogram bins, is set to 64. We tested image features: *Image, Wavelet* and *HoW* features at each level, denoted as *Wavelet-l* and *HoW-l.* Moreover, we combined all levels of *Wavelet* and *HoW*, denoted as combined *Wavelet* and combined *HoW,* and tested as well.

Table 4 shows the dimensionality of each feature with different patch sizes. *Wavelet-l* dimensionality becomes larger as the level increases, while *HoW-l* has constant dimensionality.

**Table 4.**
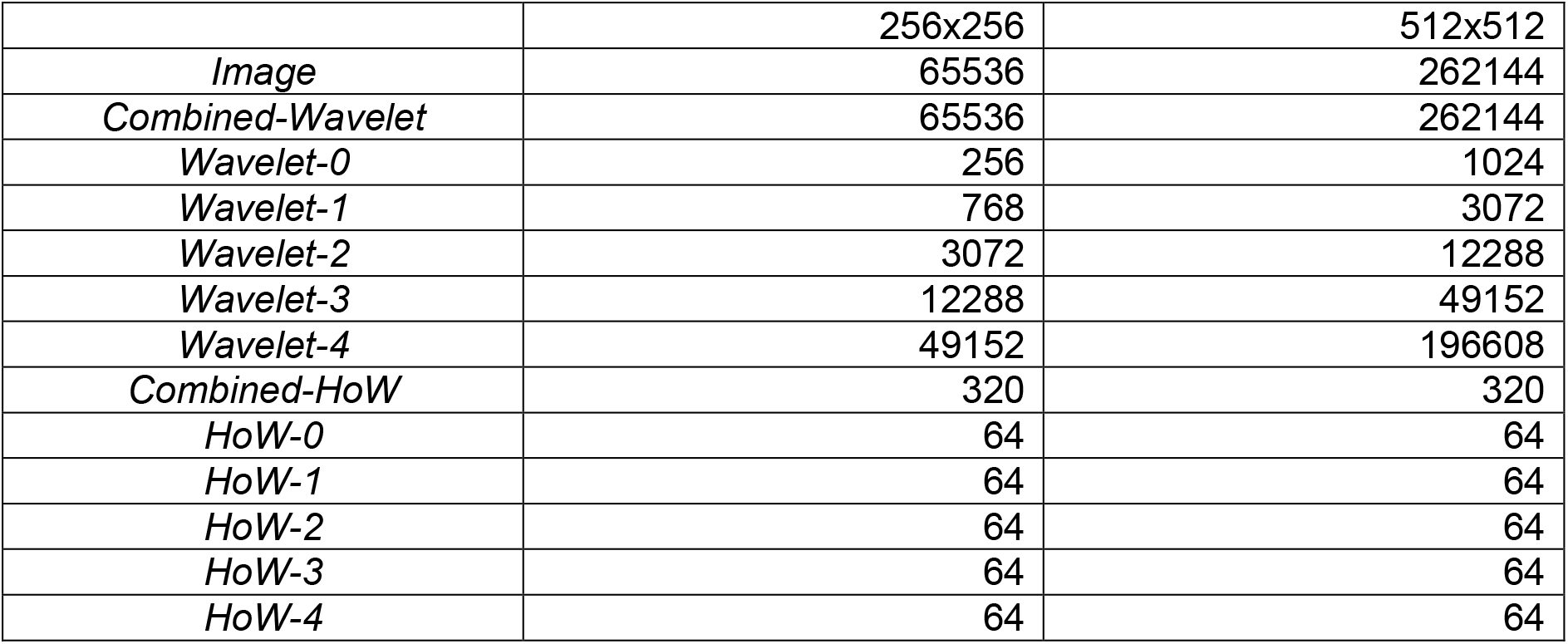
The dimensionality of the image features.

### Elapsed time of feature extraction

The first experiment measures the elapsed time of the feature extraction process. Figure 6 visualizes the elapsed time of the image processing steps. Note that the pre-processing is executed per raw image while the feature extraction is executed per patch. Thus, patch size affects the elapsed time only on the feature extraction. The pre-processing is roughly 4 Hz that is far from real-time applications that require at least 30 Hz. The feature extraction is dominated by Wavelet transformation; however total elapsed time for *HoW* computation is roughly 60 Hz even for the larger patch size.

**Fig 6.**
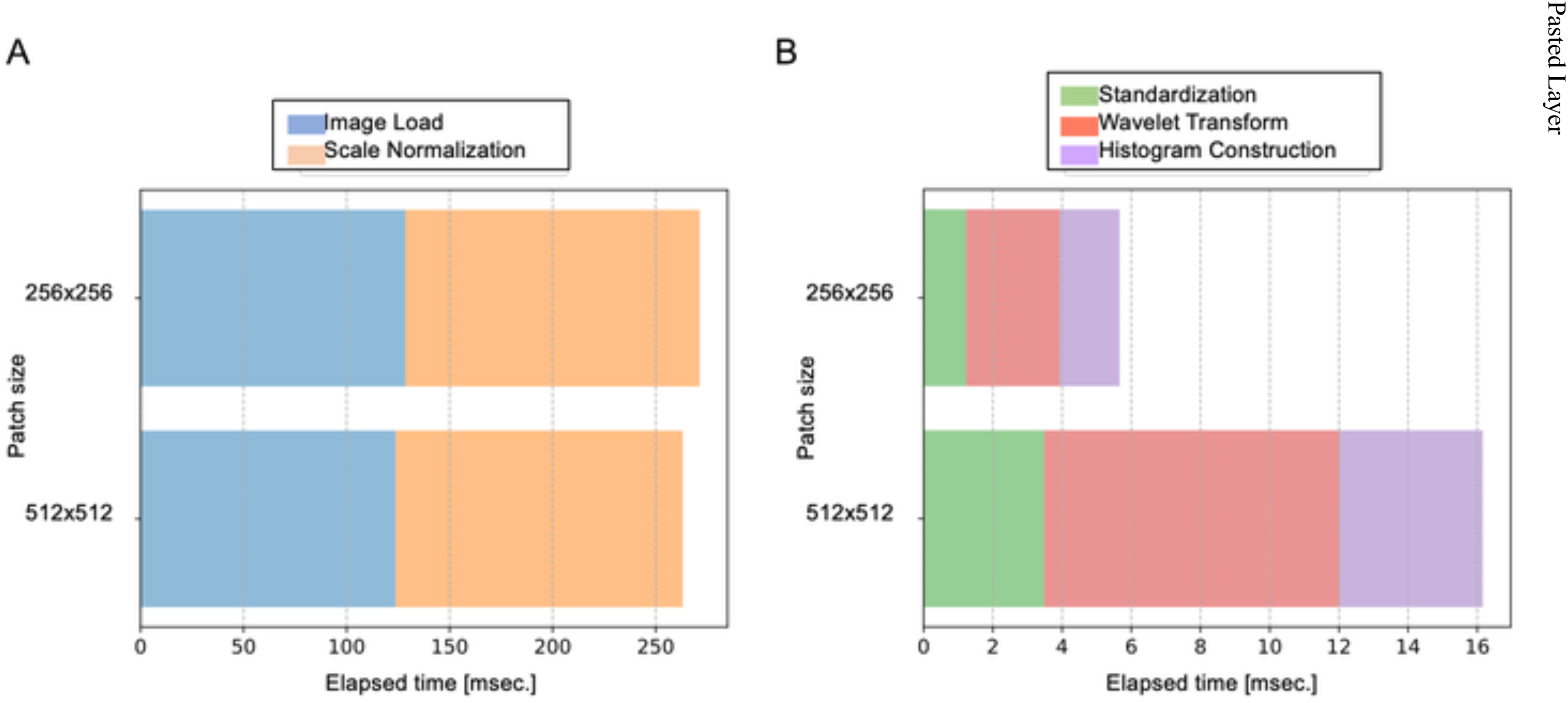
The elapsed time of the image processing steps [msec.]. (A) The elapsed time of pre-processing. (B) The elapsed time of feature extraction.

To improve the computational speed, we can use both hardware and software level techniques. The hardware technique is to use solid state drive (SSD) instead of HDD so that the process *Image Load* is fastened. One of potential software techniques is parallel programming that simultaneously processes more than a single image such as multi-core programming and graphics processing unit (GPU) programming. To improve and globalize the computing environment, a potential application can be a cloud-based system that users send raw MAT images via the internet and all the processes are performed on the server PC with multi-core CPU and GPU.

### Qualitative evaluation of MAT image feature classification

The second experiment is a qualitative evaluation of the MAT image feature classification. In this experiment, we visualize the distribution of all the data in each image feature domain to see which image feature is suitable for image classification.

For this evaluation, we use T-distributed Stochastic Neighbor Embedding (t-SNE) ^10^. In t-SNE, a non-linear dimensionality reduction method, it embeds high dimensional data into lower dimensions of two or three dimensions in such a way that similar data are distributed closer and dissimilar data are distributed further in the lower dimension with high probability.

When t-SNE embeds image features of each class into distinctly isolated clusters, the features have potential ability to be good features for classification.

We set the hyper-parameter of t-SNE as follows: perplexity is 50 and the number of iterations is 3000. We applied t-SNE visualization for all combinations of image features and resolutions of 256 × 256 and 512 × 512.

Figure 7-19 compares t-SNE visualization of each image feature. In each plot, red x and purple + symbols represent negative and positive data respectively and values of both axes are omitted because the scale does not matter in the embedded lower dimensional spaces. As shown in the figure, both negative and positive data of *Image* and *Combined-Wavelet* are embedded into inseparable clusters (Fig. 7 and 8). On the other hand, higher level *HoW*, levels 2 to 4, and all levels are embedded into distinctly isolated clusters as expected (Fig. 17-19). Lower-level *Wavelet*, levels 0 and 1, show intermediate embedding of aforementioned ones that the embedded features look inseparable with lower resolution 256 × 256 but more separable with higher resolution 512 × 512 (Fig. 9 and 10).

**Fig 7.**
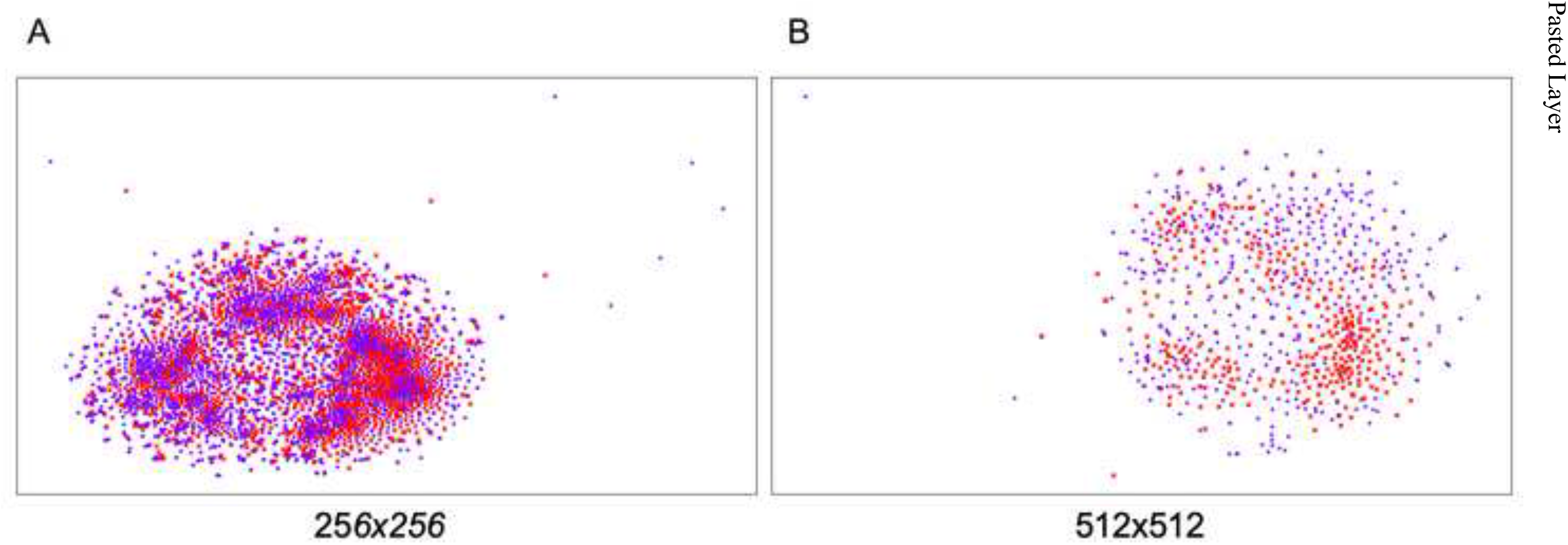
The t-SNE 2D embedding of the feature *Image*. Red x and purple + symbol represent the feature of negative and positive patches respectively.

**Fig 8.**
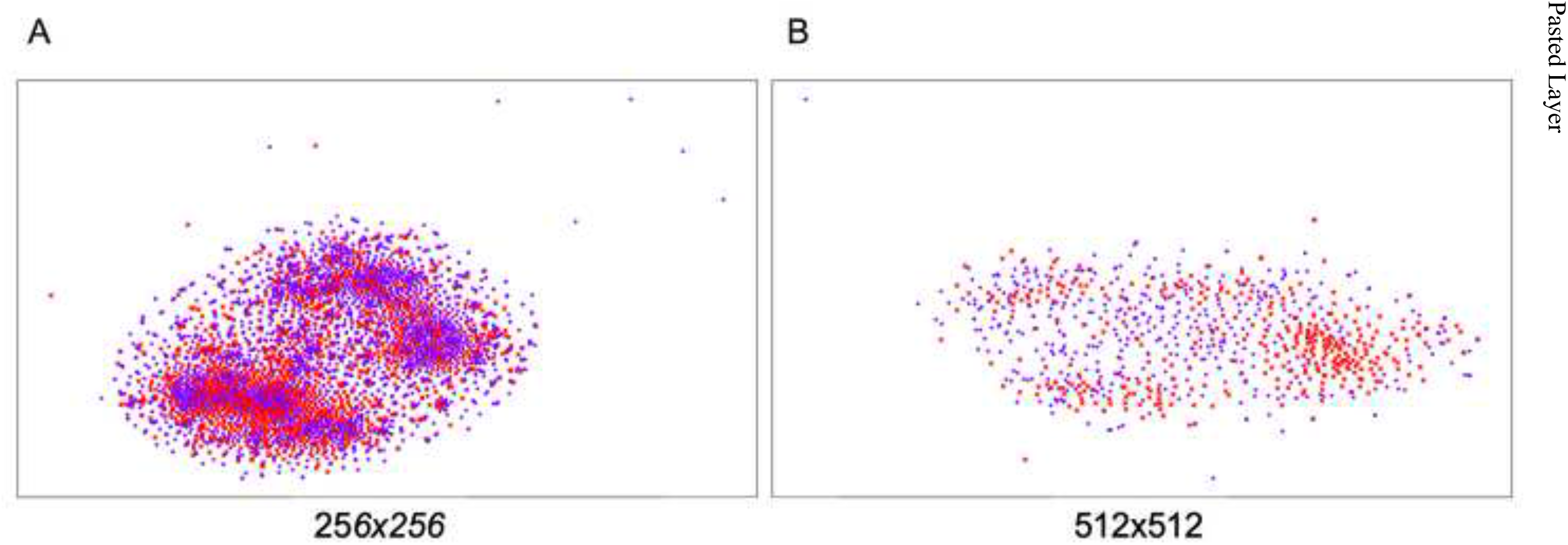
The t-SNE 2D embedding of the feature *Combined-Wavelet*. Red x and purple + symbol represent the feature of negative and positive patches respectively.

**Fig 9.**
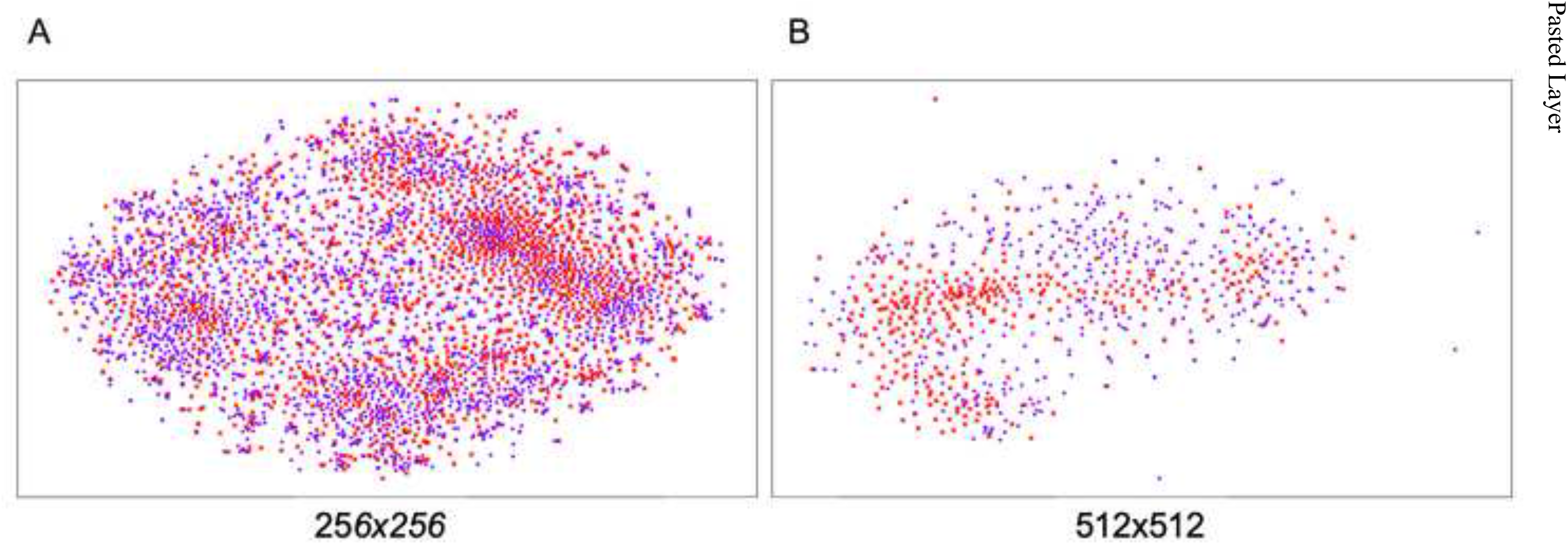
The t-SNE 2D embedding of the feature *Wavelet-0*. Red x and purple + symbol represent the feature of negative and positive patches respectively.

**Fig 10.**
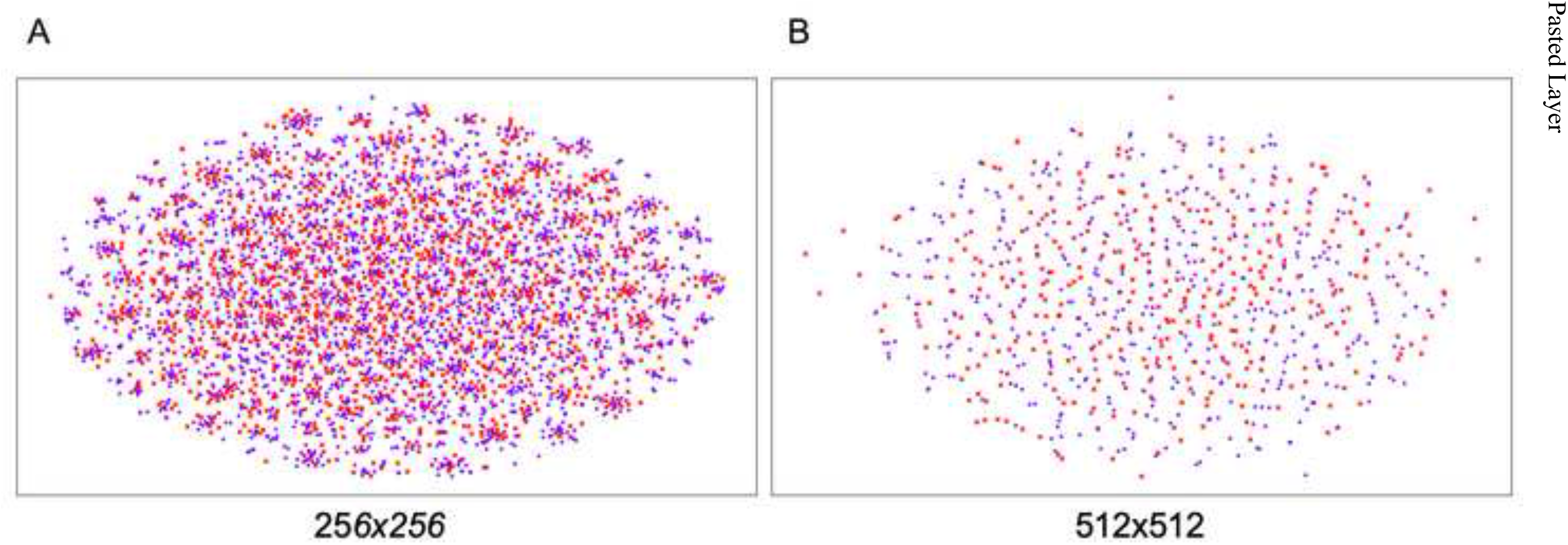
The t-SNE 2D embedding of the feature *Wavelet-1*. Red x and purple + symbol represent the feature of negative and positive patches respectively.

**Fig 11.**
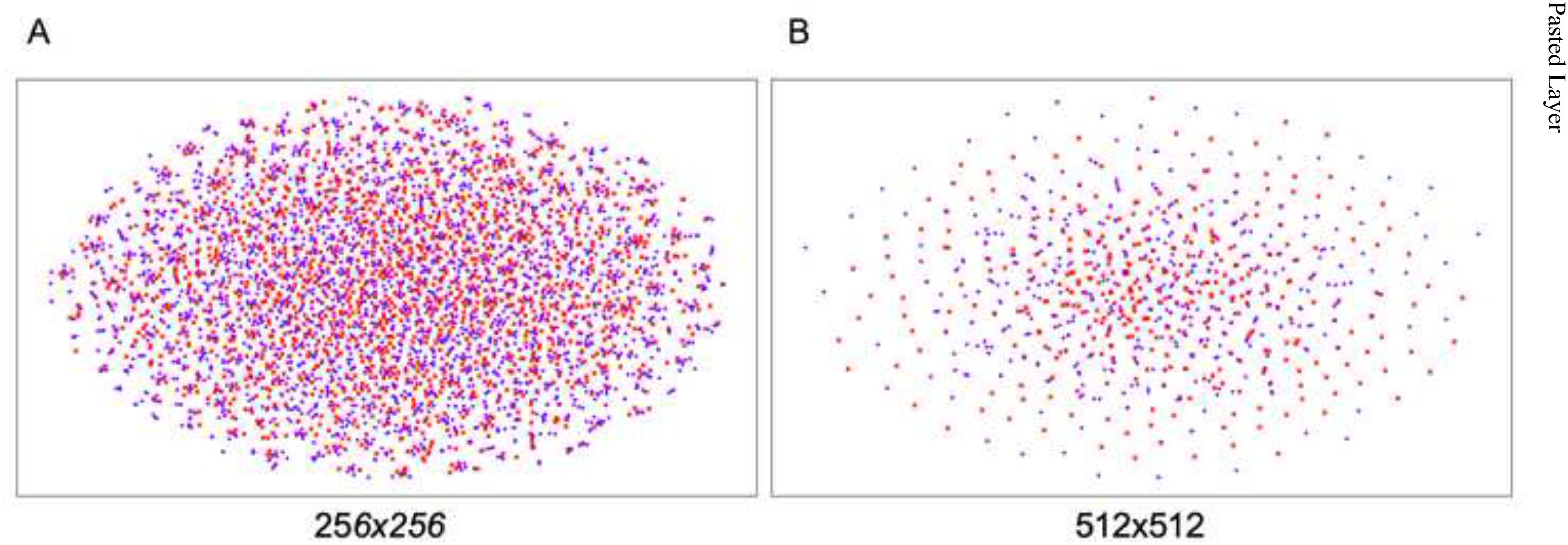
The t-SNE 2D embedding of the feature *Wavelet-2*. Red x and purple + symbol represent the feature of negative and positive patches respectively.

**Fig 12.**
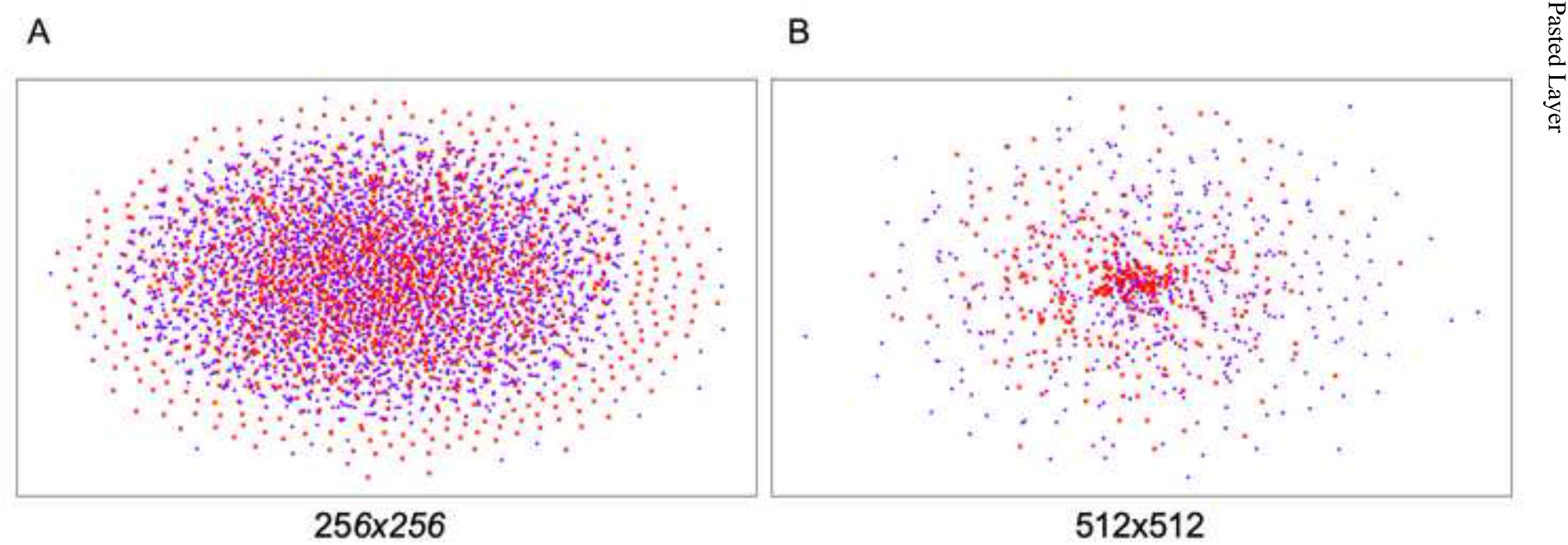
The t-SNE 2D embedding of the feature *Wavelet-3*. Red x and purple + symbol represent the feature of negative and positive patches respectively.

**Fig 13.**
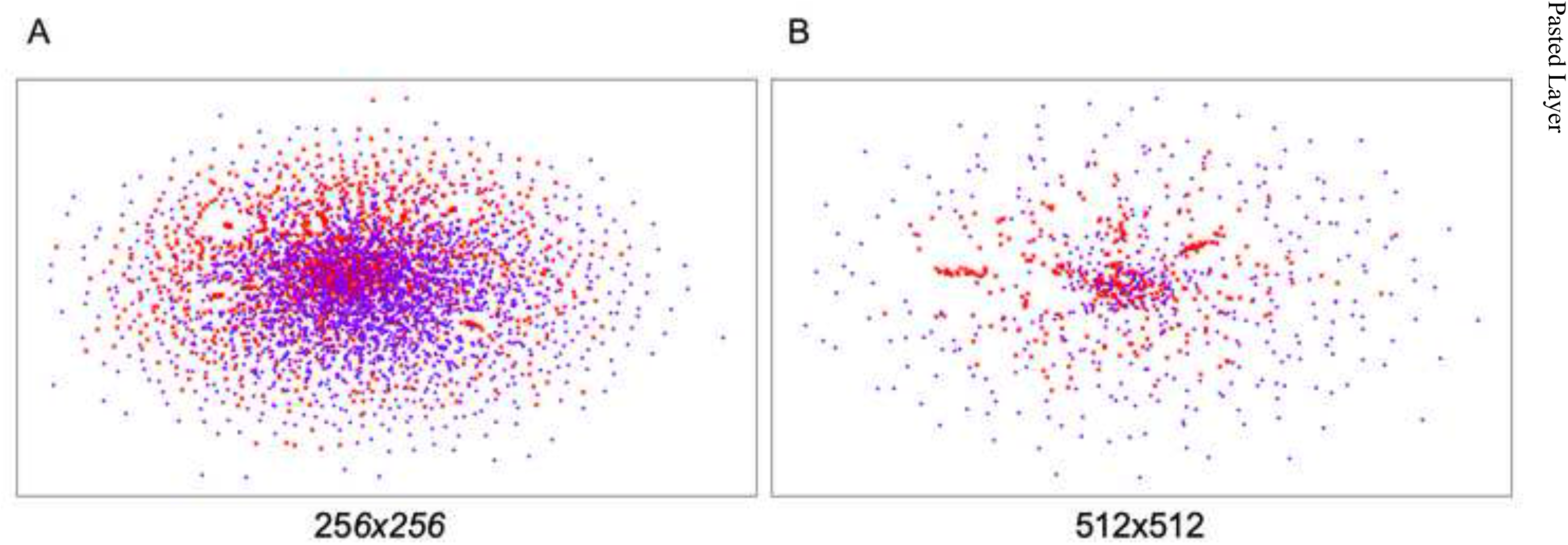
The t-SNE 2D embedding of the feature *Wavelet-4*. Red x and purple + symbol represent the feature of negative and positive patches respectively.

**Fig 14.**
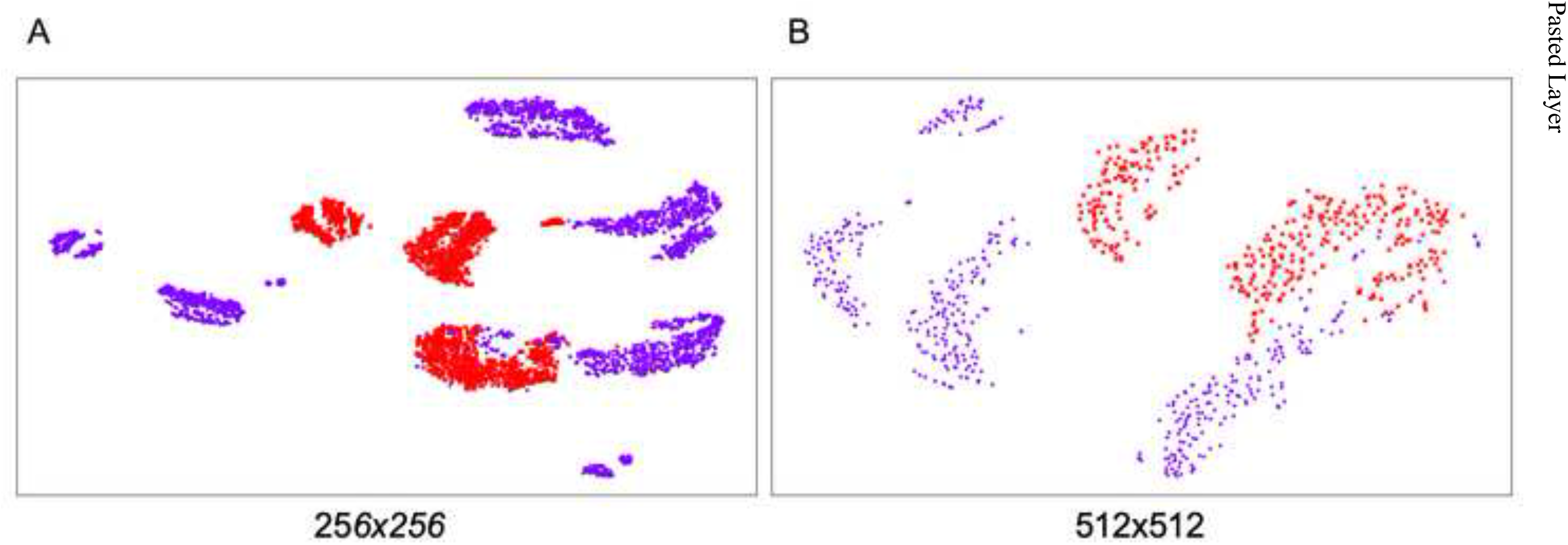
The t-SNE 2D embedding of the feature *Combined HoW*. Red x and purple + symbol represent the feature of negative and positive patches respectively.

**Fig 15.**
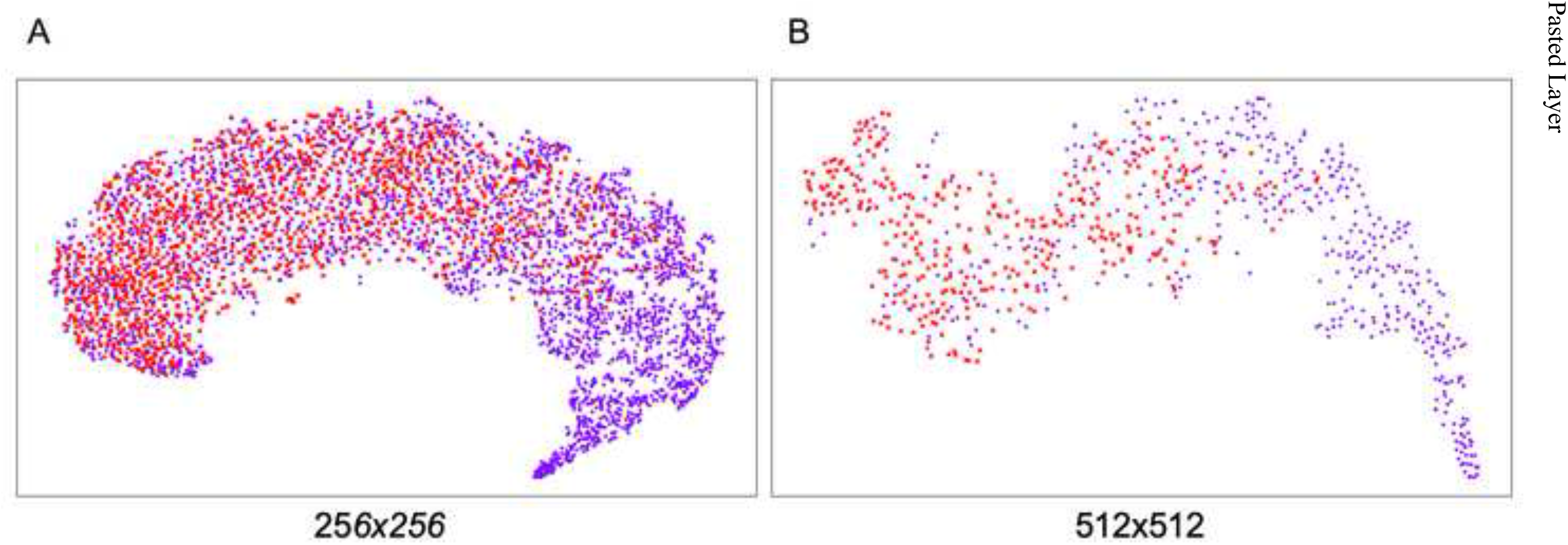
The t-SNE 2D embedding of the feature *HoW-0*. Red x and purple + symbol represent the feature of negative and positive patches respectively.

**Fig 16.**
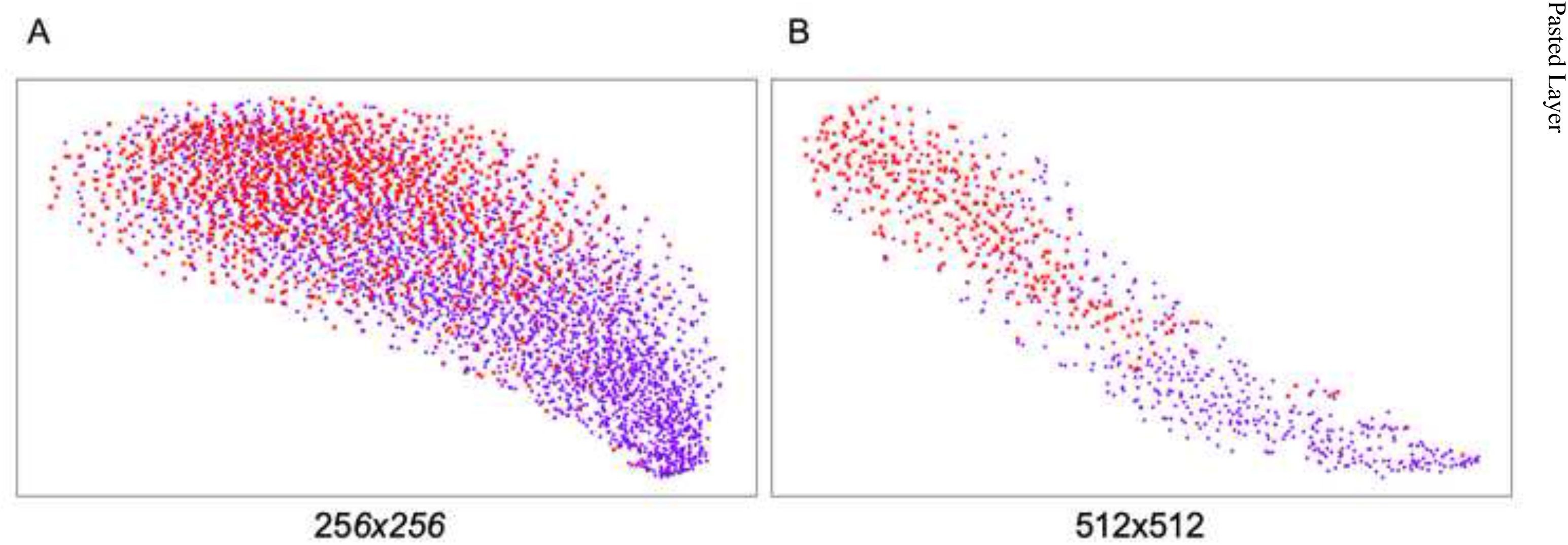
The t-SNE 2D embedding of the feature *HoW-1*. Red x and purple + symbol represent the feature of negative and positive patches respectively.

**Fig 17.**
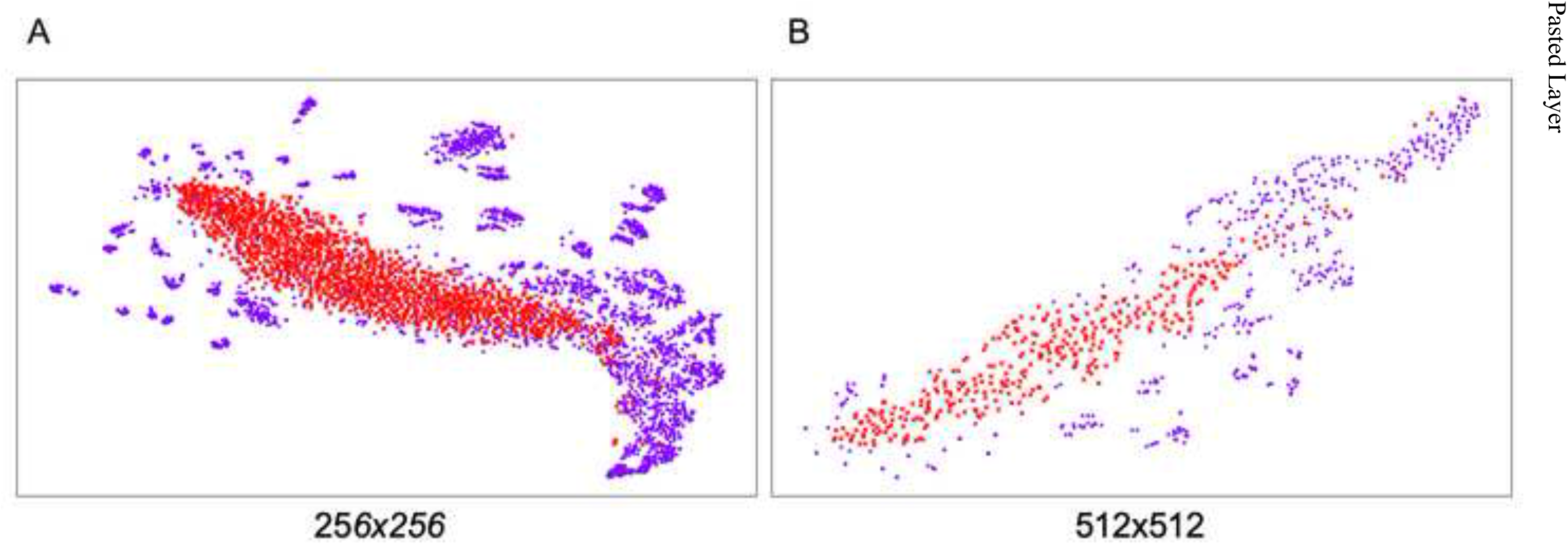
The t-SNE 2D embedding of the feature *HoW-2*. Red x and purple + symbol represent the feature of negative and positive patches respectively.

**Fig 18.**
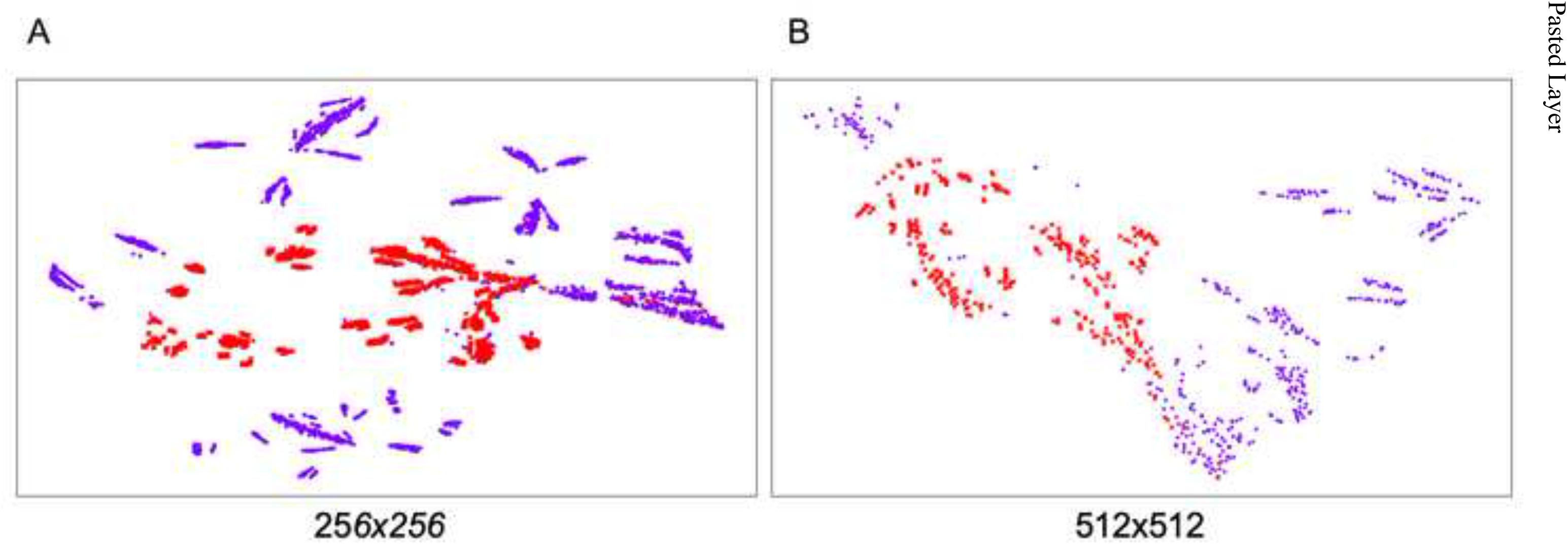
The t-SNE 2D embedding of the feature *HoW-3*. Red x and purple + symbol represent the feature of negative and positive patches respectively.

**Fig 19.**
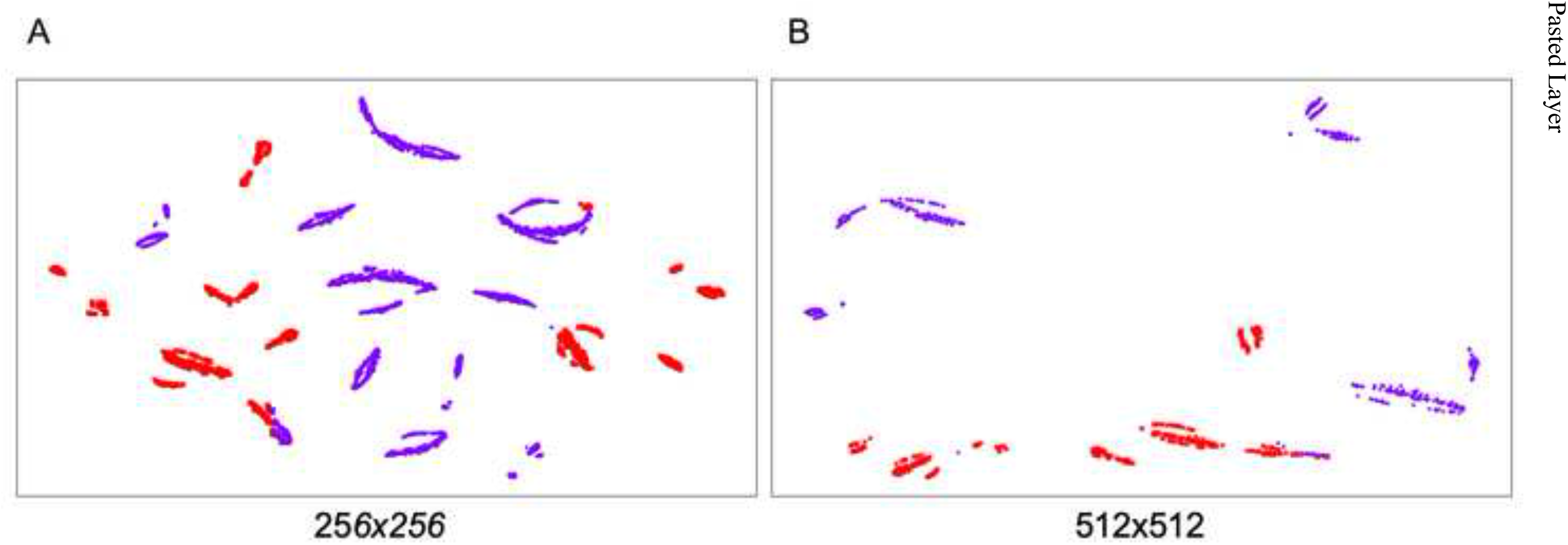
The t-SNE 2D embedding of the feature *HoW-4*. Red x and purple + symbol represent the feature of negative and positive patches respectively.

For further investigation, Figure 20 compares *HoW* features of the training data with patch size 256 × 256. The shape of *HoW*-0 varies even in the same category while *HoW*-4 seems to have a unique shape for each category. This indicates that SVM well-classifies higher level *HoW*s but mis-diagnoses lower level *HoW*s.

**Fig 20.**
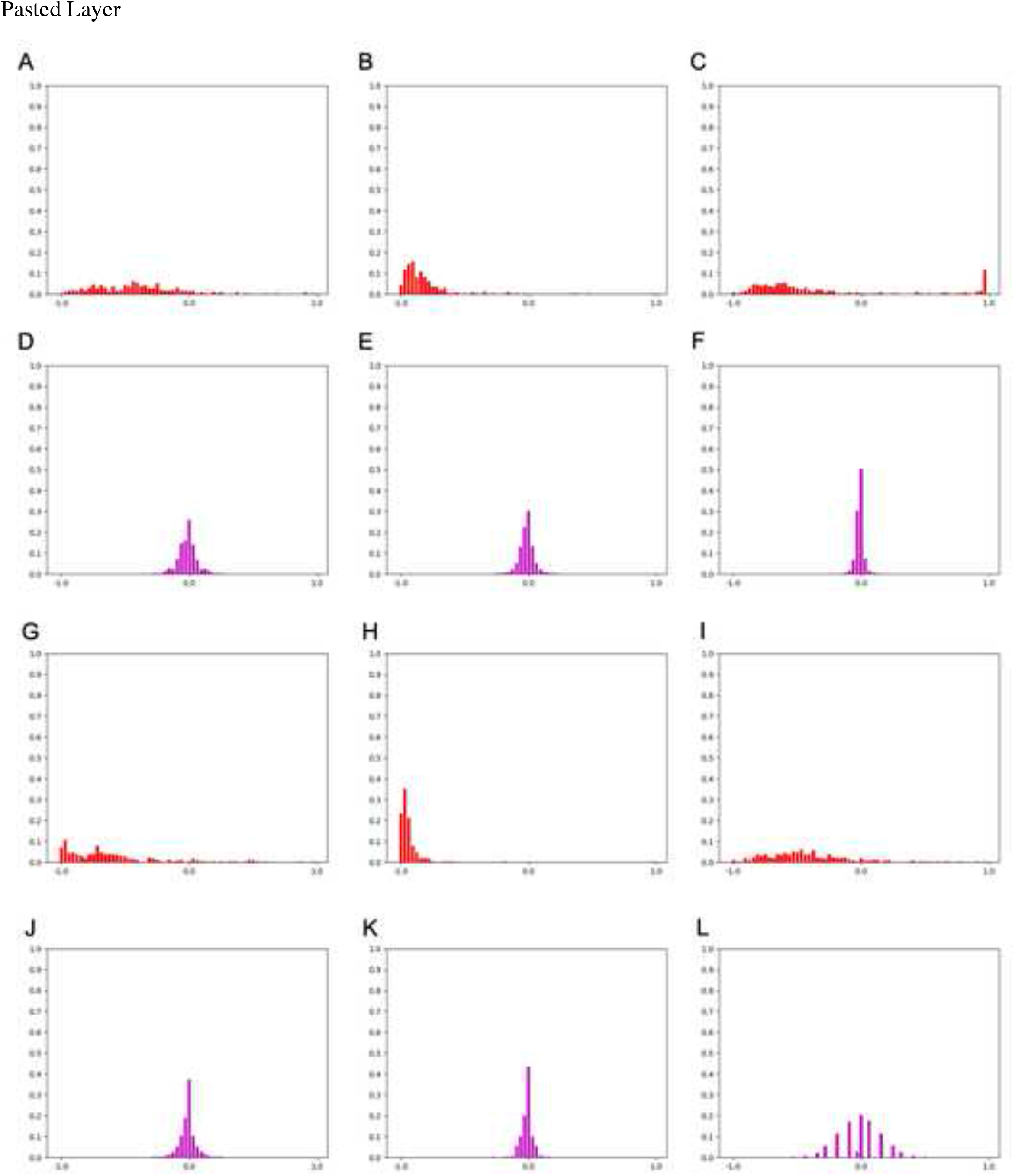
A comparison of *HoW-0* and *HoW-4* features of patch size 256 × 256. (A)-(C) *HoW-0* features of Negative data. (D)-(F) *HoW-4* features of Negative data. (G)-(I) *HoW-0* features of Positive data. (J)-(L) *HoW-4* features of Positive data.

### Quantitative evaluation of MAT image feature classification

The third experiment is a quantitative evaluation of the MAT image feature classification. In this experiment, we conducted an image classification experiment with K-fold cross validation to quantitatively evaluate the image features.

For SVM, we use radial basis function as kernel and potential hyperparameters which are described as below:

*C* ∈ {10^*i*^ | *i* = 0, 1, 2, 3} and *γ* ∈ {10^*i*^ | *i* = −3, −2, −1, 0}.

For K-fold cross validation, we use 60% of data for training and the remaining 40 % for test data. The number of folds *K* is set to 5. As evaluation criteria, we use Matthews Correlation Coefficient (MCC) ^11^) that ranges from -1 to 1, -1, where -1 means all predictions are wrong, 0 means the predictions are equivalent to random prediction, and 1 means all predictions are correct. Contrary to F-measure, more frequently used criteria, MCC considers balance ratios of two classes that is crucial for our case.

Figure 21 shows confusion matrices and MCC values of each image feature with different patch sizes. Definitions of graph keys are as follows: True Negative, negative data is predicted as negative; False Positive, negative data is predicted as positive; False Negative, positive data is predicted as negative; True Positive, positive data is predicted as positive. In the figure, horizontal bars with \ symbol represent true prediction while ones with / symbol do false prediction. In both resolutions, higher-level *HoWs* result in higher MCC while lower-level *HoW*, Image, and *Wavelet* result in lower MCC.

**Fig 21.**
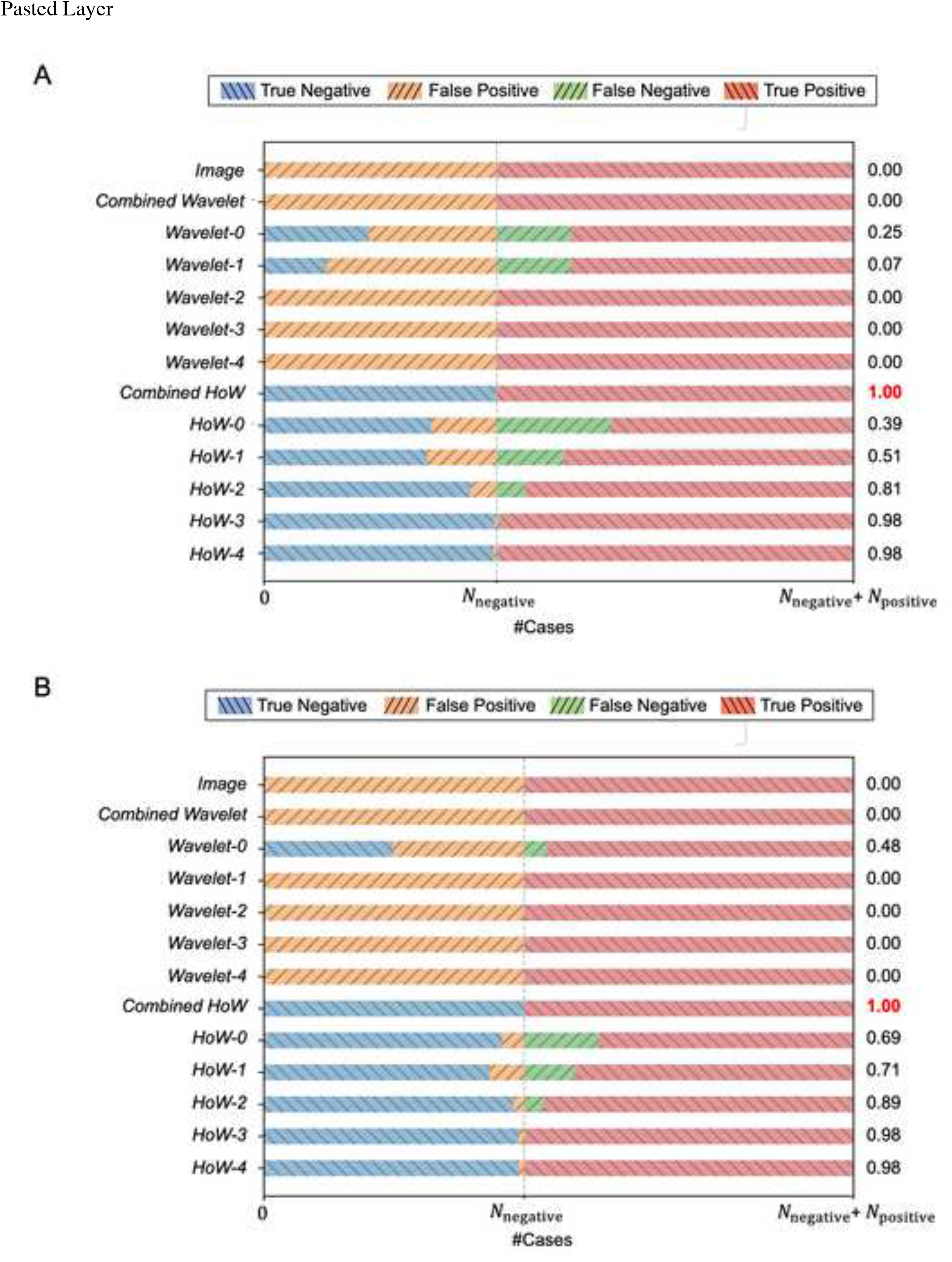
Visualized confusion matrix and MCC for each feature of different patch sizes. (A) Patch size 256 × 256. (B) Patch size 512 × 512.

Figure 22 shows failed test cases of *HoW* features. Comparing those failed features to the training data shown in Figure 9, it is convincing that the SVM mis-classified those features. Table 5 and 6 show the elapsed time for the training and test respectively. Note that the elapsed time of the test is measured per each patch while the one of the training is measured per all the training dataset. Comparing the data between patch sizes, the elapsed time of 256 × 256 cases is 10 times and 5 timers larger than 512 × 512 cases, on training and test cases respectively. Considering the elapsed time and classification performance, we can say that HoW efficiently codes the characteristics of MAT images. This result indicates that SVM with higher-level *HoWs* work as a good MAT image classifier.

**Fig 22.**
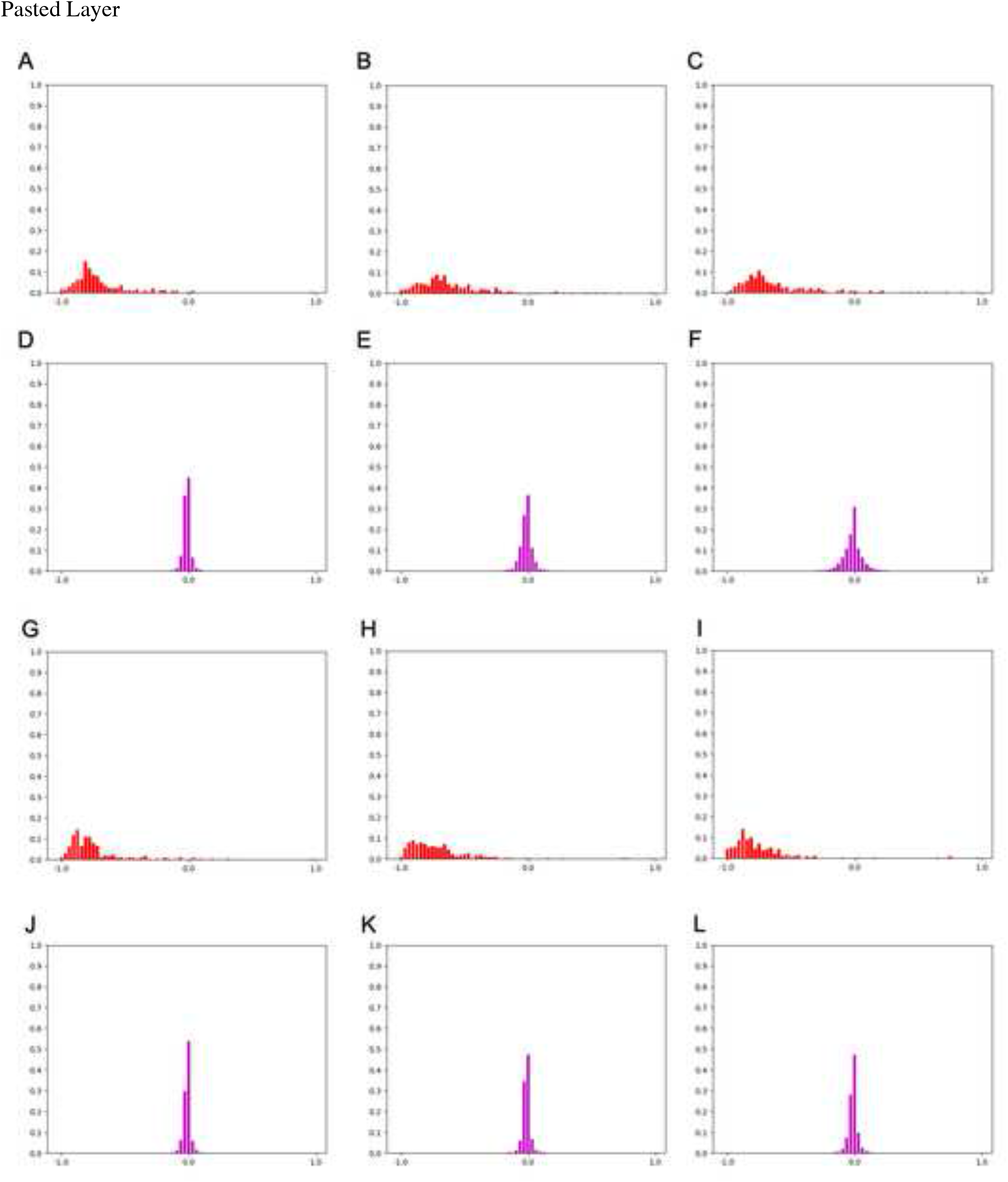
A comparison of *HoW-0* and *HoW-4* features from the failed cases of patch size 256 × 256. (A)-(C) *HoW-0* features of false negative cases. (D)-(F) *HoW-4* features of false negative cases. (G)-(I) *HoW-0* features of false positive cases. (J)-(L) *HoW-4* features of false positive cases.

## Discussion

MAT is a standard test for diagnosis of leptospirosis and a determination training is required for clinical technologists, though there have still been variations among technicians or facilities. An alternative to MAT is the macroscopic slide agglutination test (MSAT), developed by Galton et al.^12^. The MSAT is considered a rapid, practical, easy and accessible test, and is as sensitive as the MAT ^13^. The MSAT was originally developed for the serological diagnosis of leptospirosis in humans, mainly for the screening of acute and recent cases of infection ^14^. The MSAT suggests the possible infective serovars using an antigen in suspension, which may include a pool of up to three inactivated serovars. The MSAT can be used for the diagnosis of leptospirosis in both humans ^15, 16^ and animals ^17–19^.

Other serological assays, such as the enzyme-linked immunosorbent assay (ELISA) ^20, 21^ can also be used to detect infection. However, these alternatives are just simple rapid screening tests, and MAT is needed as ever in high-density areas where precisely testing with more serovars. We aim to automate and standardize MAT, the conventional and standard diagnostic method, rather than developing a new method to replace MAT. We propose to introduce a machine learning approach for determination of antibody titer.

In this study, we primarily attempted to standardize MAT by automating the determination process. Our experimental results showed that the proposed machine learning-based pipeline with the derived image feature well-recognized the agglutination in MAT images, especially combined *HoW* resulted in the MCC score of 1.0, which means perfect prediction. This indicates that the proposed method could substitute the skilled examiners’ knowledge and experience by using machine learning techniques. As we showed in the current paper, adding parameters progress capability of analysis, further tuning of the image feature may improve the classification performance.

There still remains some space to improve the proposed method. The standard MAT examination procedure by comparing MAT images to a reference image, while the proposed method classifies each MAT image based on the amount of agglutination areas within. To refine the proposed method, we need to establish an appropriate way to compare reference and MAT images in image feature space. We plan to apply several histogram comparison methods and select one with the best classification performance. Or we will design another image feature that shows a clear difference between positive and negative data when we compare reference and MAT images.

Although a lot of image data was acquired as a result, the limitation of this study is that we tested only one serotype condition, serovar Manilae, with a positive and a negative serum, and an image acquisition device. We need to study and validate with data from other serovars and devices but conclude at least that the direction of the image analysis has been adequately investigated.

A potential future direction is to directly tackle the situation even with smaller numbers of data. One of the recent trends in the machine learning community focuses on algorithms working with less data. To name a few, semi-supervised learning and transfer learning are good candidates for our problem. Semi-supervised learning algorithms are designed for situations that small amounts of samples are known for their positivity or negativity. Transfer learning algorithms alter knowledge that was built with a large amount of labeled data for a specific problem to another problem where we have access to a small amount of labeled data.

The current study suggests that MAT will be fully automated in the future. Usually, MAT requires to transfer samples from 96-well plates to glass slides, and this is the most complicated handling in all processes. However, if we automate all the manual procedures, it is possible to develop a quick MAT method.

We picture the final product of the proposed method as a cloud-based system. By launching this kind of system on the cloud, anyone can utilize the system via the internet and therefore we can support people in poor resource situations. Furthermore, we can collect a huge amount of data from all over the world that is an image of the future research style.

## Conclusion

This paper aimed to build a machine learning model on MAT as our first step toward ultimate goal to automate the MAT procedures for the diagnosis of Leptospirosis. Our idea was to introduce a typical machine learning-based image classification pipeline that represents images by an appropriate feature and uses an SVM to classify each MAT image based on the difference in the feature space. The conducted experiments validated that *HoW* feature is an efficient and effective feature for MAT image classification. From these evidence, we concluded that the machine learning-based image classification pipeline has a potential power of fully automated MAT which we are in process of developing.

## Acknowledgements

We are grateful to Mr. Shoji Tokunaga for his advice.

## Author Contributions

Conceptualization; Y.O., R.O. Data curation; Y.O., R.O.

Formal analysis; Y.O., R.O., T.M., S.M., Y.N., M.S., S.Y.A.M.V.

Funding acquisition; Y.O., R.O., J.F. Investigation; Y.O., R.O. Methodology; Y.O., R.O.

Project administration; R.O.

Resources; T.M., S.M., Y.N., M.S., S.Y.A.M.V.

Software; Y.O. Supervision; J.F.

Validation; Y.O., R.O., T.M., S.M., Y.N., F.O., M.S., S.Y.A.M.V., J.F.

Visualization; Y.O.

Writing - original draft; Y.O., R.O.

Writing - review & editing; Y.O., R.O., T.M., S.M., Y.N., F.O., M.S., S.Y.A.M.V., J.F.

## Funding

This work was partially supported by JSPS KAKENHI Grant Number 18K16174 to R.O., the discretionary fund of Tottori University President to Y.O. and R.O, and the Research Program of the International Platform for Dryland Research and Education, Tottori University to J.F.

## Notes

### Competing Interest Statement

The authors have declared no competing interest.

